# PhageTransformer for scalable and accurate bacteriophage host predictions

**DOI:** 10.64898/2026.08.29.748026

**Authors:** Malte Siemers, José L. López, Bas E. Dutilh

## Abstract

Bacteriophages can only be understood through their interactions with bacterial hosts. As environmental sequencing efforts expanded, the number of available phage genome sequences has exploded, yet the vast majority of these sequences lack host information. Predicting the host of a newly observed phage is therefore a key challenge in virology.

Several computational tools can predict phage-host relationships from genomic data, but they share notable limitations: (1) the number of different hosts that can be predicted remains relatively restricted; (2) tools tend to assign confident host predictions to non-viral input sequences; and (3) most tools have a trade-off between accuracy and speed.

Here we present PhageTransformer (PT), a deep learning model for phage-host prediction that addresses these limitations. We benchmark PT against existing tools on 3,881 independent phage-host pairs from GenBank and public HiC data, and demonstrate that it achieves competitive or superior prediction accuracy at greatly reduced runtime.

## Introduction

Bacteriophages depend on their hosts in nearly all stages of their life cycle, and understanding their biology requires linking them to the bacteria they infect. Experimentally establishing this link mostly requires propagating the virus on cultured bacteria, restricting the approach to cultivable organisms and interactions that can occur under laboratory conditions^1,2^. Meanwhile, the rapid expansion of environmental sequencing has generated viral genomic data at an increasing pace over the past two decades^3^, producing vast collections of metagenomically derived virus genomes and genomic fragments for which no host information is available. This discrepancy necessitates computational approaches that can generalize from limited experimental knowledge to the diverse and poorly characterized environmental virosphere.

Host prediction is complicated by the fact that phage-host specificity often varies at the strain level, governed by rapidly evolving molecular determinants. Phage-host adsorption depends on specific molecular interactions between phage receptor-binding proteins and host surface structures, which are among the most variable elements in both phage and host genomes^4,5^. Additional strain-level variability arises from the gain and loss of defense and anti-defense systems, yielding interaction patterns that vary between closely related strains^6^. Host-range commonly extends to the genus rank, suggesting that if the receptor binding protein (RBP)-receptor interaction and defense-anti-defense systems are aligned, infection could occur. A higher-level modular structure is also observed in phage-host interaction matrices, although those are not always taxonomically resolved^7,8^. As the genome sequence is generally conserved between related phages infecting the same genus^9^, these observations hold promise for computational host prediction based on sequence data^1,10^.

A range of computational methods has been developed to predict phage-host interactions. Classical approaches rely on direct sequence signals such as k-mer similarity between phage and host genomes^11^, CRISPR spacer matching^12^, or sequence homology^13^. Increasingly, machine learning and deep learning models have been applied to phage-host prediction (Table 1). Convolutional neural networks have been trained on encoded genome representations to classify hosts directly from nucleotide sequences^14,15^. Other approaches operate at the protein level, classifying hosts based on protein family profiles derived from phage proteomes^16–18^ or structure-aware embeddings of receptor-binding proteins^19^. Integrative methods combine multiple signals, either by fusing nucleotide and protein features within a single model^20^, unifying predictions from existing tools through a meta-classifier^10^, or constructing multimodal knowledge graphs^21^. A recent benchmark of 27 tools confirmed that no single method is universally optimal, revealing a critical trade-off between prediction accuracy, sensitivity, and computational cost^22^.

**Table 1.** Machine learning-based phage-host prediction tools. The number of hosts is listed at the indicated prediction rank. GCN: graph convolutional network; MLP: multilayer perceptron; CNN: convolutional neural network; DT: decision tree; RBP: receptor-binding protein.

| Tool | Year | Method | Prediction rank | Number of hosts | Training set size | Tested on |
| --- | --- | --- | --- | --- | --- | --- |
| RaFAH<sup>16</sup> | 2021 | Random Forest on Proteome | Genus | 617 | 25,879 | 3,053 phages (190 genera) |
| CHERRY<sup>21</sup> | 2022 | Multi-Modal GCN | Species | 233, extendable | 1,260 | 615 pairs (88 species) |
| vHULK<sup>17</sup> | 2022 | MLP on Proteome | Genus & Species | 77 & 118 | 8,616 | 2,153 phages (77 genera) |
| DeepHost<sup>14</sup> | 2022 | CNN on Genome | Genus & Species | 72 & 118 | 7,880 | Random 10% subset |
| PHERI<sup>18</sup> | 2023 | DT Classifier on Proteome | Genus | 50 | 4,723 | Random 20% subset |
| iPHoP<sup>10</sup> | 2023 | Integrative | Genus | Not reported, | 17,105 | 1,870 phages |
|  |  | Framework |  | extendable |  | (170 genera) |
| PHIStruct<sup>19</sup> | 2024 | MLP on RBP embedding | Genus | 7 | 7,627 | ESKAPEE (7 genera) |
| CoMPHI<sup>20</sup> | 2025 | Composite Machine Learning | Genus | 256 | 3,018 | Random 30% subset |
| PhageCGR Net<sup>15</sup> | 2026 | CNN on Genome | Genus & Species | 72/118 & 108/206 | ~8,756 | Random 20% subset |
| PhageTransformer (PT) | This study | CNN + Transformer on Genome | Genus | 1,084 | 104,620 | Stratified 20% subset (1084 genera) |

One important factor underlying these trade-offs is how each method constructs a numerical representation of the phage genome. Some predict and encode proteins^16–19^, while others encode the full genome, for example as a two-dimensional Chaos Game Representation^15^ or as a matrix of one-hot encoded nucleotides^14^. K-mer-based approaches decompose sequences into short words of fixed length, each mapped to a numerical representation, which can be informed by known biochemical properties of the underlying nucleotide or amino acid composition^20^. Alternatively, input feature (token) representations can be learned directly from data, as in deep learning models for natural language processing and protein sequence analysis^23–28^.

While such learned embeddings have not yet been adopted in phage-host prediction tools, they allow the representations to be optimized jointly with the model architecture. Two components are particularly relevant in this context. Convolutional neural networks (CNN) learn short sequence motifs and combinations thereof by sliding learned filters along an input^29^. Applying a convolutional layer to a tokenized sequence produces a per-position profile of all learned features, while a stride reduces the number of positions in the output, effectively compressing information along the sequence. Stacking multiple convolutional layers into a convolutional tower yields increasingly abstract, hierarchical combinations of sequence motifs at progressively larger scales. Transformers complement this architecture by placing features in context^30^. The self-attention mechanism central to the transformer architecture learns the importance of both individual and combined features, enabling signal integration across the full input length^25–27,31,32^.

While CNNs have been applied to phage-host prediction^14,15^, the combination of learned embeddings, convolutional feature extraction, and transformer-based integration has not yet been explored in this context. Current phage-host training datasets, typically comprising fewer than 10,000 phage-host pairs from a few hundred host genera (Table 1), limit both the taxonomic scope of the predictor and the choice of model architectures. Moreover, prediction performance is generally reported as a single aggregate metric over an unstratified subset of the target host categories, making it difficult to assess whether model performance is dominated by a few well-represented taxa.

Here, we present PhageTransformer (PT), a general-purpose deep learning model for phage-host prediction across 1,084 host genera directly from nucleotide sequences. We tokenize phage sequences into consecutive, non-overlapping 3-mers simultaneously in all six reading frames (3 forward, 3 reverse, see Figure 1A), capturing both nucleotide composition and protein-level information in a single representation, invariant of the input strand orientation. An end-to-end model architecture^31,32^ combines a strided convolutional tower that extracts local sequence features with two hierarchical transformers that integrate the local patterns with overall genomic structure. This enables PT to learn sequence-level signals predictive of host range, such as similarities in codon usage and purine content between phages and hosts^33,34^, as well as host-specific sequence motifs in both inter-and intragenic regions^1,10^. PT is freely available at https://github.com/MGXlab/phagetransformer.

**Figure 1.**
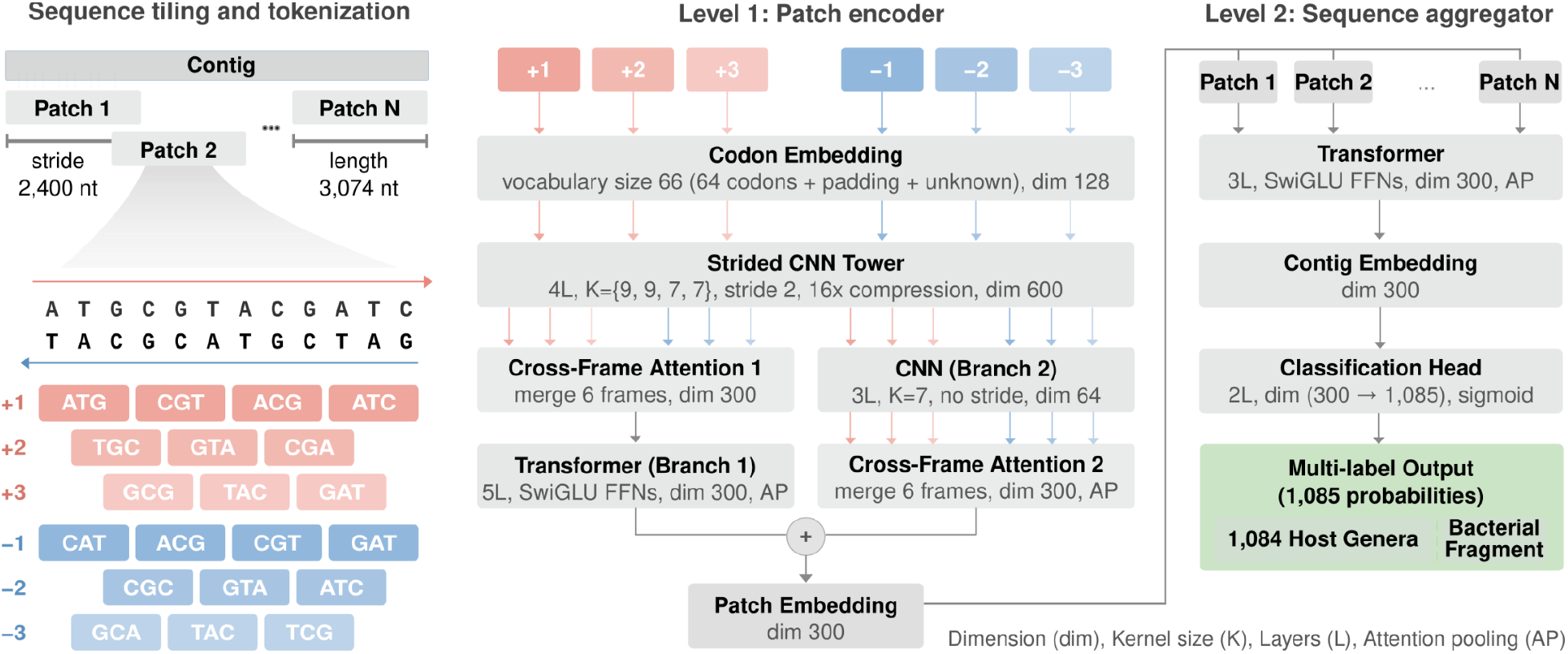
PhageTransformer architecture. Input DNA is tokenized into trinucleotides across six reading frames, processed by a strided CNN tower and cross-frame attention to produce patch embeddings, then aggregated across the full genome by a second transformer with attention pooling. A dense classifier outputs independent sigmoid predictions for 1,084 host genera and a bacterial fragment class.

## Methods

### Construction of a phage-host interaction dataset with 1,084 host genera

Training the PT model required two labelled datasets: a dataset of phage genomes labelled with their host genus, and a dataset of genus-labeled host genomes for the contamination-detection class, that also let the model learn shared compositional signatures. Because the model was trained as a supervised multi-label classifier over host genera plus a bacterial fragment class, every sequence in both datasets carried a genus label from a shared label space, with bacterial sequences additionally labeled as bacterial fragments.

To obtain a broad range of phage-host pairs, we assembled the phage genome dataset from multiple sources, as described previously^35^. Briefly, we combined prophages extracted from GenBank^36^ bacterial genomes using PhiSpy^37^ (https://doi.org/10.25451/flinders.22317059), phage sequences from PHD^38^ (accessed February 12, 2025), IMG/VR4^39^ (IMG_VR_2022-09-20_7.1), and the prophage subset of Prophage-DB^40^ derived from GTDB^41^ (https://datadryad.org/dataset/doi:10.5061/dryad.3n5tb2rs5). Sequences were retained only if host taxonomy was reported to at least the genus rank (n=1,837,034/5,687,759). Redundancy was reduced by using Vclust 1.3.0 at ≥95% nucleotide identity over ≥85% of the shorter contig^42^ (approximately species level), and representative sequences for each cluster chosen as the sequence at the mode of the length distribution. We subsequently quality-assessed all sequences with geNomad^43^ (v1.11.2) and CheckV^44^ (v1.0.1 with database v1.5), trimmed identified host-flanking regions, and discarded sequences if more than 20% of their predicted genes were CheckV-assigned host genes. This yielded an intermediate dataset of 204,150 representative dereplicated phage sequences, annotated with 10,049 different host genera (Table S1).

To further reduce redundancy while preserving diversity of viruses and hosts, we constructed a protein-sharing network among all viral sequences. We predicted CDS using pyrodigal^45^ (v3.6.3, meta=True, closed=True), and clustered translated protein sequences with diamond^46^ (v2.1.8, --approx-id 90, --member-cover 90). For each sequence, we computed the mean number of other sequences sharing each of its genes, providing a per-genome measure of local density within the dataset. This measure, together with the annotation frequency of host genera, was used to derive a probability per phage genome to be included in the final training set (p∼1/(ln(n sequences per host genus)*(mean number of phages sharing each protein))), applying sparse sampling to phages infecting highly represented host genera and denser sampling to those infecting underrepresented ones. Host genera represented by fewer than 12 phage sequences after sampling were discarded. The resulting phage genome dataset comprised 104,620 sequences annotated with 1,084 different host genera (Table S1).

The host genome dataset enabled training PT to both identify prokaryotic contamination and leverage shared nucleotide signatures and frequencies between phages and their hosts. It comprised 16,039 GTDB r226 species representative genomes from the 1,084 bacterial host genera represented in the final phage genome dataset (Table S1). To mask prophage regions and minimize the overlap between phage and host genome sequences, we aligned our phage genome dataset against the host genome dataset with minimap2^47^ (v2.26) and masked all regions in host genomes with alignments longer than 2,000 nt or 20% of the corresponding phage genome (Table S1).

### Architecture of the PhageTransformer model

PhageTransformer processes DNA sequences hierarchically at two levels (Figure 1A). On level 1, input nucleotide sequences are first divided into overlapping patches of fixed length (default size: 3,072 nt, stride: 2,400 nt). For each patch, trinucleotide sequences in all six reading frames (three forward, three reverse complement) are extracted and passed through a patch encoder consisting of a codon embedding layer (66 wide for 64 codons, 1 padding token and 1 token for unknown/ambiguous, dimension 128), a convolutional neural network (CNN, 4 layers, kernel sizes {9,9,7,7}, stride 2, 600 channels), cross-frame attention to exchange information between reading frames, and a transformer (5 layers, SwiGLU FFNs, dimension 300, attention pooling) that produces a single embedding per patch. A lightweight convolutional branch (3 layers, kernel size 7, no stride, 64 channels, attention pooling, projection to dimension 300) operates in parallel, and its output gets added to the output of the transformer to generate a final patch embedding. A sequence-level aggregator transformer (3 layers, SwiGLU FFNs, dimension 300, attention pooling) then attends over all patch embeddings of the strided input sequence and produces multi-label host genus predictions via an attention-pooled classification head (2 dense layers, internal dimension 300, output dimension 1,085) followed by a sigmoid function. The rest of the model employs GELU activation functions. The final output dimension corresponds to the 1,084 possible host genera plus one bacterial fragment class to capture bacterial sequences expected in metagenomic input data. In this configuration, the model had 19,070,337 learnable weights. For details on the choice of individual parameters see Supplementary Note 1.

### Training procedure

We trained the model in two successive phases to first learn host-specific features from local context (within 3,074 nt) and subsequently aggregate local signals across a contig or genome. In the first phase, we pre-trained the patch encoder using a temporary classification head on individual patches of 3,074 nt length. We employed an oversampling strategy that extracted more patches per epoch from sequences of rare host genera (up to 3-fold higher coverage than abundant host genera) to balance the long-tailed distribution, enabling predictions for rare genera with improved confidence. To train the model to detect host contamination and leverage shared sequence information between phages and their hosts, bacterial genome patches were included at a ratio of 10% relative to the phage patches, labeled with both the “bacterial fragment” class and the corresponding host genus. In the second phase, we froze the pre-trained patch encoder and trained the sequence-level aggregator on full phage genomes, again supplementing each epoch with 10% independently sampled bacterial genome fragments with a length distribution corresponding to the phage sequences (3,606-199,912 nt, mean: 35,639 nt). Because diversity differs strongly between host genera (n=2-517 species), the decision which genome a sequence got sampled from was based on a frequency proportional to a power of the number of species per genus (α=0.25), yielding an approximately four-fold difference in coverage per genome between the most and least diverse genera. Both phases used focal binary cross-entropy loss (gamma=1.8) with class-frequency-based positive weights (proportional to the inverse of the class frequency, capped between 2 and 6) to mitigate label imbalance. To strengthen the model’s ability to reject sequences without any host signal, 10% of training sequences were randomly shuffled and received a zero-label. After training, we applied post-hoc temperature scaling on the validation set to calibrate output scores to more closely reflect class probabilities (multiplied with raw model logits before sigmoid scaling), and determined score thresholds at controlled false discovery rates (10% and 20%).

### Model evaluation

We trained three model configurations. Two of them, the “validation model” and the “phage-only model”, were trained on a subset of the available data so that they could be evaluated on the held-out remainder. The third, the “production model”, was trained on all available data.

To investigate the effect of adding host genome sequences to the training of PT, we trained one model on sequences derived from the phage and host genome datasets (validation model) and one without bacterial sequences and the bacterial fragment class (phage-only model). For both of these models, the phage genome dataset was divided using a genus-stratified 80/20 train-validation split. For the validation model, we also added a single host genome per genus at random from the host genome dataset. Restricting training to one genome per genus prevented overlap between validation regions in one host genome and training regions of closely related species within the same genus. Each of these genomes was then partitioned by selecting one contiguous region spanning 20% of the total genome length at random for validation, with the remaining sequence used for training. The production model was trained on the complete phage genome dataset and on all 16,039 host genomes at full length, without held-out regions. We used the validation model to compute all validation metrics for phage-host prediction, bacterial fragment detection, and taxonomic classification across the 1,084 host genera (Figure 3, Figures S2-S5), the phage-only model to show host scores within bacterial genomes (Figure S3), and the production model for the comparison against other host-prediction tools on independent datasets (Figure 4, Figure S6).

To evaluate model predictions across all 1084 host genera, we report precision, recall, micro-, and macro-averaged F1 scores, and area under the precision-recall curve (AUPRC). Precision measures the proportion of correct predictions among all predictions for a given class; recall measures the proportion of correctly predicted test cases among all true cases for that class. F1 combines these two measures in a geometric mean and serves as a joint indicator for accuracy and sensitivity of the predictor. The four measures are defined as:

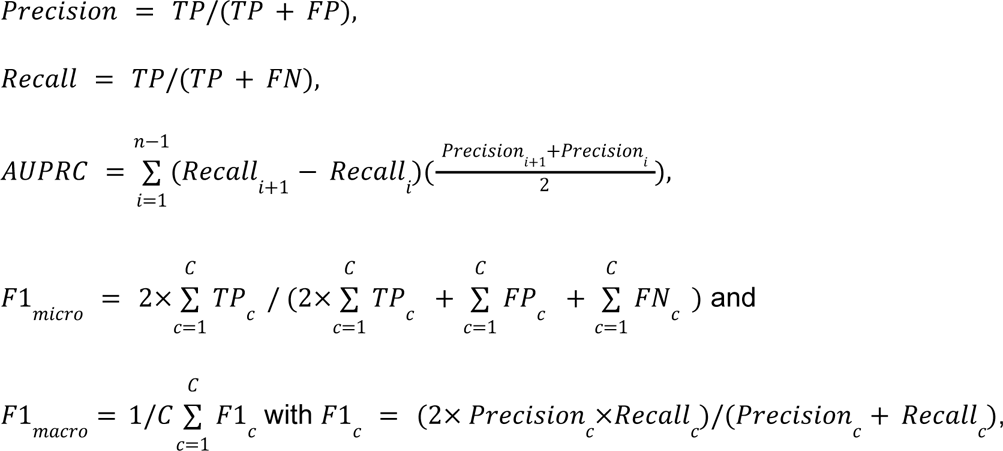

where C is the number of possible classes, FP is the number of false positives, TP is the number of true positives, and FN the number of false negatives. AUPRC were computed on n=200 evenly spaced score cut-offs from 0 to 1.

Because these prediction metrics are only defined relative to a decision threshold, we calibrated one. We first computed the tANI between the phage sequences used in validation and in training with Vclust (1.3.0, align command). We stratified the validation phages by their maximum tANI to any training phage and used the most distant stratum (tANI_max_≤40%) to set the threshold at which the false discovery rate reached 10%, giving a score cutoff of 0.69 for host prediction classes and 0.5 for the bacterial fragment class. These cutoffs were applied wherever a thresholded metric is reported.

### Comparison to other host-prediction tools and taxonomy standardization

To benchmark PT against existing tools, we composed three datasets of phage-host pairs that were not present in the training data of any of the tools: (1) GenBank^36^ phages with annotated hosts submitted after June 2025 (training or reference data of all compared tools was collected until then or earlier), referenced as the “GenBank dataset”, (2) a collection of quality-controlled HiC-derived datasets from a recent study^48^, referenced as the “HiC viral dataset” and (3) three additional HiC-derived datasets^49–51^, together referenced as the “HiC contaminated dataset”. For (1), we used the Entrez module of biopython to download sequences from NCBI Nucleotide using the query string “txid2731619[Organism:exp] AND 2025/07/01:3000[PDAT] AND srcdb_genbank[PROP]” and filtered the resulting set for sequences with a host annotation (n=1,269/2,400). For (2), we downloaded the Supplementary Data of a recent ecological study^48^ (https://doi.org/10.5281/zenodo.14851637). We filtered the MGE contigs using the annotated geNomad viral score and host annotation (genomad_viral_score≥0.7, host assigned to genus or species rank, n=2,612/6,572). The remaining sequences had nine different biomes annotated. For (3), we downloaded the HiC evaluation dataset published in a recent benchmarking paper^22^ (https://doi.org/10.5281/zenodo.16975470, n=251). Then, geNomad (v1.11.2) was used to classify sequences into viral, chromosomal and plasmid if their corresponding score was above 0.7. Sequences without any score above 0.7 were classified as “other” (Table S2).

As the host labels in these datasets reference either NCBI taxonomy or various outdated versions of GTDB taxonomy^41^, we mapped all host labels to GTDB release 226 using our own tool gtdb-translate (r226, https://github.com/maltesie/gtdb-translate), to ensure consistent evaluation across datasets and tools. Translations from NCBI taxonomy were based on the mode of the NCBI-to-GTDB correspondences derived from the GTDB 226 metadata file. For forward-mapping across GTDB versions, the tool constructs a directed acyclic graph of renaming and reclassification events from the taxdump files maintained at https://github.com/shenwei356/gtdb-taxdump, and traverses this graph to find the most probable current label for any taxon name present only in older versions.

We installed iPHoP (v1.4.1, with database June 2025) and CHERRY (within phabox2 v2.3.1, with database v2.1 from June 2025) and ran both with default parameters without adding custom reference genomes. We forward-mapped iPHoP’s GTDB output and translated CHERRY’s NCBI taxonomy to GTDB 226 (available GTDB output was missing for multiple predictions), so that all tools were evaluated against a single, consistent taxonomy (Table S2). Of note, the implementation of CHERRY in phabox2 derives host predictions from MCL clustering in a gene-sharing network of query phage sequences and a set of reference sequences with annotated host information and not a graph convolutional neural network as reported in the original CHERRY publication and Table 1.

### Visualization of model attention weights

To interpret the PT predictions on individual phage genomes, we visualized the importance of each reading frame to the host prediction by extracting attention weights from the cross-frame attention layers (Crossframe Attention 1 and 2, see Figure 1), yielding a six-channel signal (one per reading frame) along the genome. Since frame selection happens within the patch encoder, we ran two non-overlapping tiling passes offset by half a patch length and merged them via element-wise maximum to avoid boundary artifacts. Similarly, we extracted the attention weights of the positional attention pooling layers (AP in Branch 1 and Branch 2, see Figure 1) and the patch attention pooling layer (AP in Sequence aggregator transformer, see Figure 1). The resulting weight profiles were plotted as heatmaps with coding regions predicted by Pyrodigal (v3.6.3, meta=True, closed=True) indicated as colored strips above the attention weights. All positional information is reported per CNN-compressed feature, corresponding to 16 trinucleotide tokens or 48 nucleotides in the input sequence. Overlap between the high-attention regions and annotated genes was quantified as the fraction of compressed positions at which CDS annotation and maximum attention fall into the same frame.

### On the use of generative AI

We used generative AI to write parts of the code used to train and evaluate PT, to structure and draft parts of the text of this manuscript, and to draft the design of Figure 1. We reviewed and tested all generated code, and reviewed and edited all generated text. Generative AI was not used to interpret data or to derive any conclusions presented here.

## Results

### PhageTransformer is a hierarchical DNA deep-learning model

We developed PhageTransformer, a hierarchical deep-learning architecture that predicts the bacterial hosts of phages directly from raw nucleotide sequences. The architecture processes input DNA in two stages: a patch-level encoder that extracts features from local sequence windows, and a sequence-level aggregator that combines all patch representations into a contig-or genome-wide prediction (Figure 1).

At the input layer, each patch is tokenized into trinucleotides across all possible reading frames (three forward and three reverse), yielding six parallel sequences of length 1,024 per patch (Figure 1, left). These sequences are mapped through a shared embedding layer into learned 128-dimensional trinucleotide representations. A strided CNN tower then compresses each frame along its length, trading nucleotide-level resolution for representations of local sequence motifs. Next, the model processes the compressed features in two parallel branches: In the first branch, a cross-attention merges the six reading frames per compressed position and then a transformer self-attends to the combined features, enabling the detection of complex inter-motif relationships within one patch. In parallel, a second CNN processes the compressed features in each reading frame followed by a cross-attention merge of the six frames. Attention pooling mechanisms at the end of each branch reduce their respective features to a single representation each, and both get summed into one final vector per patch (Figure 1, center). Patch embeddings of all windows in an input sequence are processed by a second transformer that models inter-patch context, followed by another attention pooling step that produces a single embedding for the query. A final dense classifier maps this representation to 1,084 host genera plus a binary bacterial detection class, using independent sigmoid activations that allow multi-label prediction, such as multiple hosts, or bacterial contamination with a taxonomic label (Figure 1, right).

### Model representations reveal biological features of the phage genome

PT’s codon embedding layer learned a representation for each of the 64 codons without explicit biological supervision. In a UMAP projection, codons encoding the same amino acid did not form compact groups, but tended to cluster together: the silhouette score of amino acid identity was close to zero but significantly higher than expected under permutation (silhouette=−0.011, p≤10⁻³; Figure 2A), indicating that the model distributed amino acid information across many embedding dimensions rather than concentrating it in a few dominant axes. Consistent with this, no single principal component dominated the embedding space: PC1 accounted for 3.7% of the total variance and PC30 still carried 1.7% (Figure S2). Notably, the single tryptophan codon TGG laid adjacent to the stop codons TAA, TAG, and TGA in the projection (Figure 2A), an arrangement consistent with the widespread reassignment of TGA from stop to tryptophan observed in diverse phage lineages^52^. Correlating the first 30 principal components of the embedding space against 15 codon properties showed that the learned representation captured both nucleotide-level and amino acid-level information, organised along separate axes (Figure S2). Purine composition was the strongest single determinant: purine content at codon position 1 and overall purine fraction both correlate strongly with PC1 (p<0.001), and purine features account for the largest share of embedding variance among all properties tested. Amino acid physicochemical properties emerge in later components, with hydrophobicity and charge at pH 7 loading on PC4 (p<0.01 and p<0.05, respectively).

**Figure 2.**
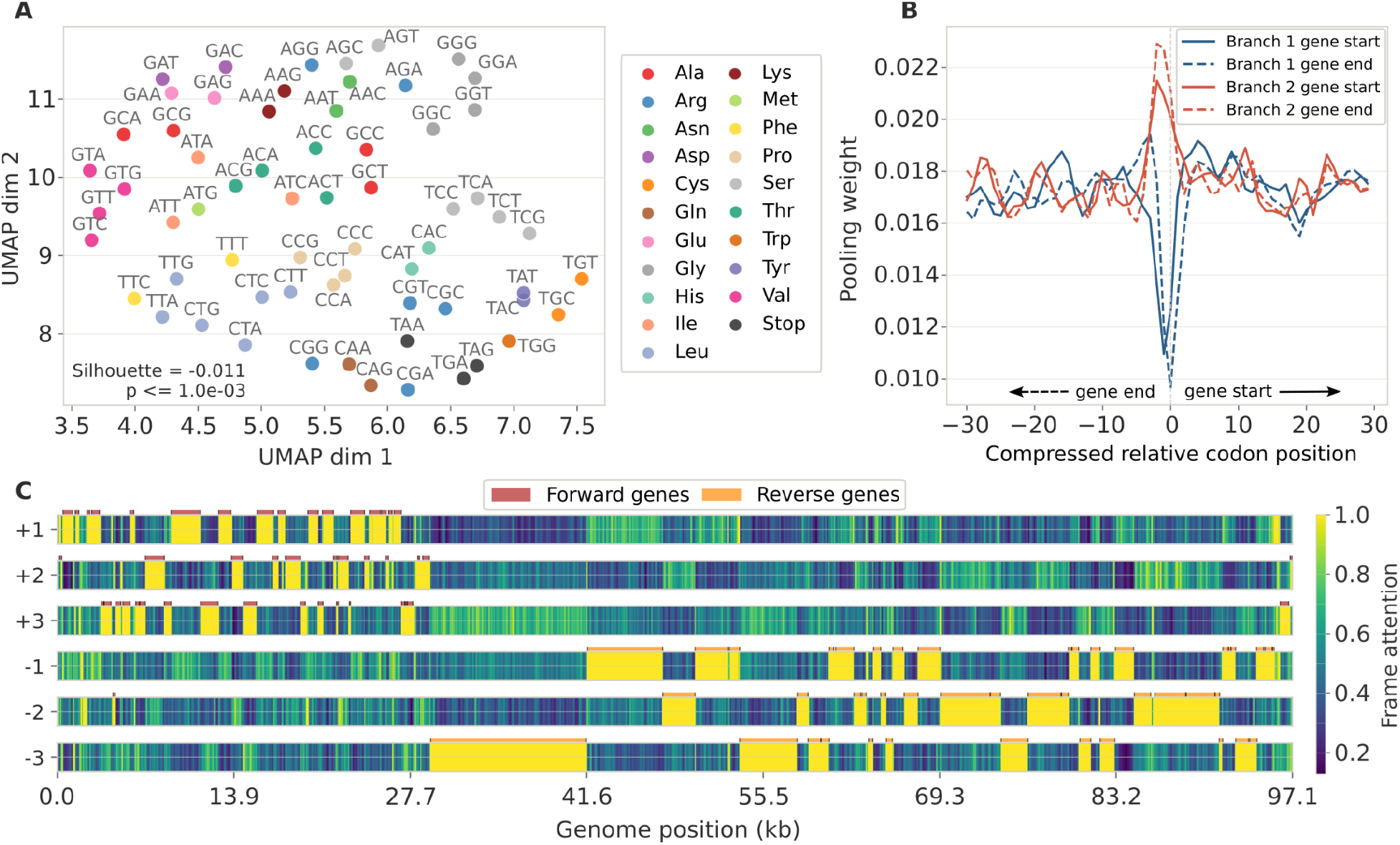
Visualization of learned representations of the genetic code structure and gene organization. **(A)** UMAP projection of the learned 128-dimensional codon embeddings, colored by the encoded amino acid. Silhouette scores (cosine distance) with permutation p-values are shown. **(B)** Average weight of the positional attention pooling per compressed (16x) codon position around gene starts (solid) and gene ends (dashed) in Branch 1 (blue) and Branch 2 (red) of the patch encoder across all CDS of 100 randomly sampled phage genomes. **(C)** Cross-frame attention weights projected onto an annotated phage genome (*Carjivirus communis* BK010471.1, 97 kb). Rows correspond to the six reading frames. High cross-attention (yellow) indicates regions that the model considers informative in this genome; the pattern tracks gene locations and strand orientation as predicted by Pyrodigal.

Next, we investigated how the weights of the various model attention mechanisms connect to biological properties of an input sequence. The two branches of the patch encoder assigned attention to different parts of a gene. Averaging positional attention pooling weights around annotated gene starts and ends revealed opposite and localised responses (Figure 2B): Branch 2 up-weighted positions at gene boundaries while Branch 1 down-weighted the same positions, with both effects confined within one or two compressed positions (the receptive field of the CNN tower covers ∼50-100 nt) of the boundary. Projection onto a full genome revealed that Branch 1 pooling weights were distributed broadly across coding regions, whereas Branch 2 weights concentrated into sharp, sparse peaks close to gene borders (Figure S3A-B). Additionally, the cross-frame attention layers of both branches recovered coding regions and their orientation. When we mapped per-frame attention weights onto a *Carjivirus communis* genome, high attention coincided with Pyrodigal-annotated genes, switching between frames in accordance with the local gene orientation (97.7% overlap, Figure 2C, Figure S3C-D). We extended this analysis to two phage genomes utilizing alternative genetic codes 4 and 15 and found attention patterns to be consistently correlated with the annotated CDS (96.7% and 91.6% overlap, Figure S4). Finally, the sequence aggregator distributed its attention unevenly across the genome (Figure S3E), indicating that not all regions contributed equally to the final prediction.

We therefore asked which genes carry the information PT relies on, by quantifying the importance of a gene as the decline in model confidence upon its removal, and testing for importance enrichment across PHROG functional categories (Figure S5). Tail-associated genes were the most strongly enriched category, at 2.5-fold over expectation (Figure S5A). At the level of individual product annotations the signal was considerably sharper: central tail fiber J and tail length tape measure protein were enriched more than 10-fold (Figure S5B). Per-genome importance distributions confirmed that these enrichments were not driven by a single gene per genome or category (Figure S5C-D). The prominence of tail fibers and tape measure proteins is consistent with their role as the primary determinants of receptor recognition and host range, indicating that PT’s predictions rest on features that mediate host specificity, rather than on genome-wide compositional similarity alone.

### Phage genomes share predictive signal with their hosts

When we applied a model trained exclusively on phage sequences (phage-only model, see Methods) to 50kb-long windows in a full *E. coli* chromosome, we found that the genus *Escherichia* was confidently predicted as a host in every window with maximum prediction probabilities between 0.44 and 0.92 (Figure S6A). This suggested that phage genomes share sufficient sequence signals with their bacterial hosts for the model to predict the taxonomy of bacterial sequences, even when trained only on phages, and that the model had picked up these compositional features.

Motivated by this observation, we extended the architecture with a dedicated class for bacterial contamination detection and included bacterial sequences in the training data (validation model and production model, see Methods). On the same *E. coli* chromosome, the validation model assigned nearly all windows to the bacterial fragment class, at higher confidence than the phage-only model achieved for its host calls (0.76-0.95), while only a small number of scattered regions retained *Escherichia* as their top label (Figure S6B).

Extending this comparison to 1,084 bacterial genomes, one per host genus in our host genome dataset, confirmed the pattern (Figure S6C). The phage-only model assigned the correct genus to a median of 0.81 of windows per genome. Under the extended validation model this rose to 0.96, whereas the fraction of windows in which the correct genus was the top label and the sequence did not get flagged as bacterial fell to a median of 0.03. All following evaluations were conducted with models including bacterial sequences in their training data.

### PhageTransformer accurately predicts bacterial hosts across taxonomic ranks

We then evaluated PT’s performance on the genus-stratified validation fraction of our phage genome dataset with the validation model (see Methods). To test the model’s ability to generalize to unseen phage genomes, we stratified the validation phage sequences by their maximum total average nucleotide identity (tANI) to any of the training genome sequences. Host prediction performance across the tANI strata declined with increasing distance from the training data, as expected, but degraded gradually rather than collapsing (Figure 3A). For phages with a close relative in the training set (tANI≥50%), 97% of genomes received a genus call above threshold, of which 96% were correct; for the most distant stratum (tANI 0-10%), this fell to 54% with 80% correct. Performance also varied substantially between taxa. At genus rank, half of all taxa were predicted with an F1 score above 0.80, rising to 0.93 at phylum rank (Figure 3B). Performance per taxon is largely a question of representation: micro-averaged AUPRC exceeded macro-averaged AUPRC at every rank (Figure S7A-B), indicating that prediction errors concentrate in host taxa that contributed few sequences. Plotting per-genus F1 scores directly against the number of corresponding training genomes confirmed this relationship (Figure S7C).

**Figure 3.**
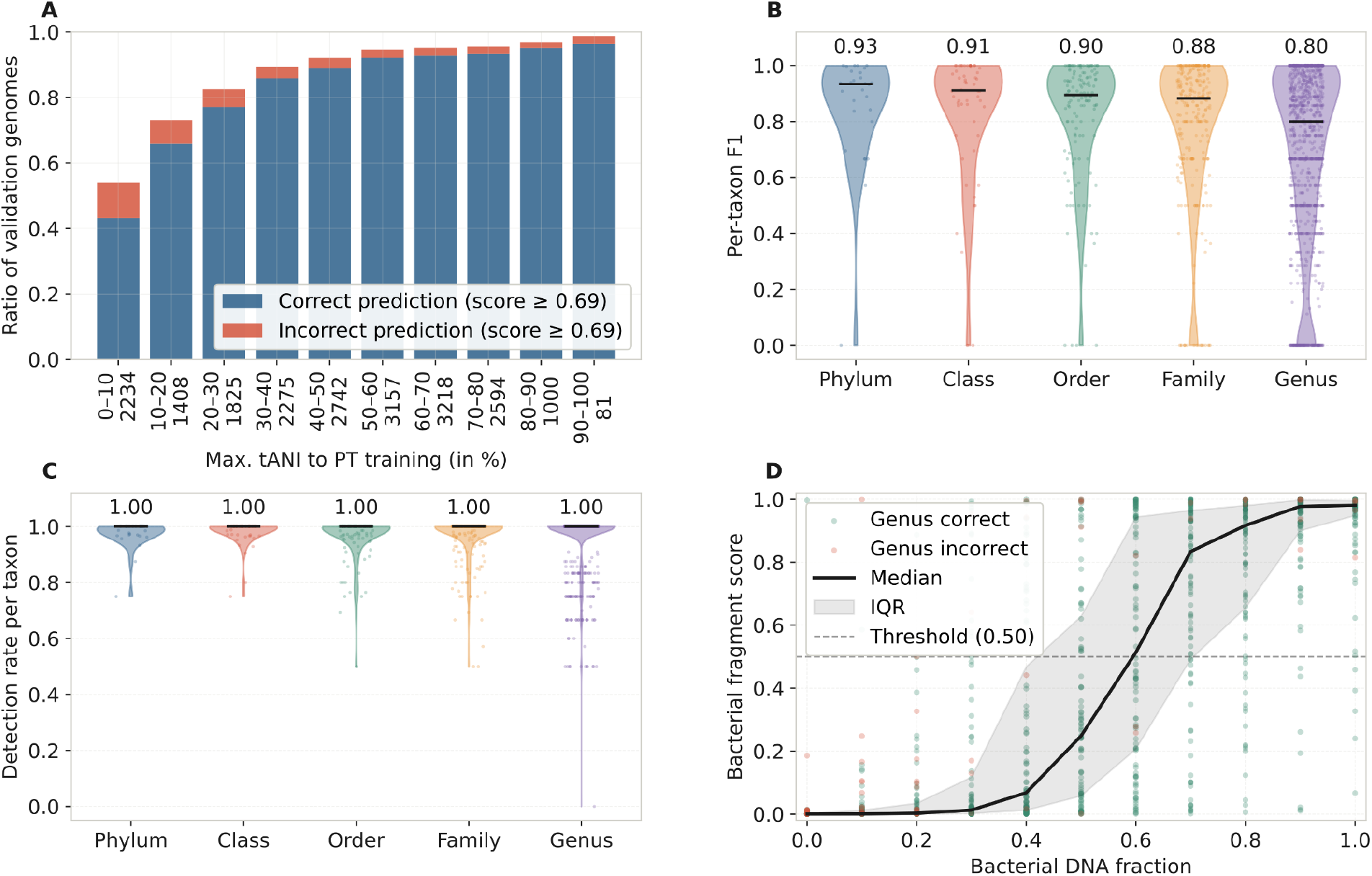
Evaluation of the PhageTransformer model on held-out validation sequences. **(A)** Prediction accuracy as a function of maximum total ANI (tANI) between the query phage and the nearest training genome. Blue: fraction of sequences with a correct prediction; red: fraction of sequences with an incorrect prediction. The number of sequences in each bin is indicated next to the bin boundaries. **(B)** Distribution of per-taxon F1 scores at each taxonomic rank. Each point represents one taxon; horizontal lines indicate the median. Violin width reflects the density of taxa at each F1 score. **(C)** Bacterial fragment detection rate at different taxonomic ranks. Each point represents one taxon at the indicated rank, violins show the distribution across 10,000 bacterial validation subsequences, and horizontal bars indicate the median. **(D)** Bacterial fragment scores in a synthetic contamination dataset. 1,000 chimeric sequences were generated from random validation phage genomes and validation region samples from their respective host genome at controlled ratios. Green dots indicate correct genus predictions for a chimera, red dots indicate an incorrect prediction. The black line and grey shadow correspond to the median and inter-quartile range (IQR) of the score distribution across phage-bacteria sequence ratios.

Next, we tested the validation model’s ability to detect and taxonomically classify bacterial sequences across 1,084 bacterial genera. The median detection rate per taxon was 1.00 at every taxonomic rank, with only a small number of genera falling below (Figure 3C). The same picture held for classification performance across taxonomic ranks, with macro-averaged AUPRC between 0.996 and 1.000 (Figure S7D) and a median per-taxon F1 of 1.00 at all ranks (Figure S7E). Since real assemblies can contain partial contamination, we tested the sensitivity of the contamination detection on chimeric sequences, constructed by combining a controlled fraction of each phage genome with DNA from its known host. The median bacterial fragment score rose monotonically with bacterial content, crossing the 0.5 threshold at approximately 60% bacterial DNA (Figure 3D). At the extremes the behaviour was well separated: 95% of sequences containing at least 90% bacterial DNA were flagged, while only 3% of sequences with 10% contamination or less were.

Finally, we tested the model’s host prediction performance on input of varied lengths by randomly drawing fragments from phage validation sequences of the 100 most abundant host genera. Sequences shorter than the patch encoder’s window size (3,074 nt) showed a sharp drop in confidence and correctness, while sequence lengths above 4,000 nt lead to a consistently high rate of correct predictions above 87% (Figure S7F).

### PhageTransformer shows competitive performance at reduced runtimes

We compared the PT production model against iPHoP and CHERRY on three independent datasets: 1,269 phage genomes in the GenBank dataset, 2,612 viral contigs spanning nine biomes in the HiC viral dataset, and 251 contigs in the HiC contaminated dataset (see Methods). Applied to the GenBank dataset, the three tools traded prediction rate against precision differently (Figure 4A). CHERRY predicted a host for 91% of sequences, with 98% of those predictions correct at phylum but only 54% at genus. iPHoP predicted for 76%, with 99% correct at phylum and 59% at genus. PT was the most conservative, predicting for 68% of sequences, but at highest accuracy across ranks: 99% correct at phylum and 80% at genus (Figure 4A). On the HiC viral contigs the ordering shifted (Figure 4B). CHERRY predicted for only 55% of sequences, with 90% correct at phylum falling to 35% at genus. PT and iPHoP behaved almost identically: PT predicted for 69% with 99% correct at phylum and 68% at genus, iPHoP for 73% with 99% and 66% correct, respectively. Because the HiC contigs were derived from nine distinct environments, we split this comparison by biome (Figure S8, Table S2). Performance was uneven across biomes for all three tools with best results reported for human-associated biomes (Figures S8A and C), and no tool dominated everywhere. PT and iPHoP recovered comparable amounts of genera at comparable precision and recall across most biomes (Figures S8A-I), while CHERRY recovered the fewest genera with lowest precision in every biome except one (Figure S8E).

**Figure 4.**
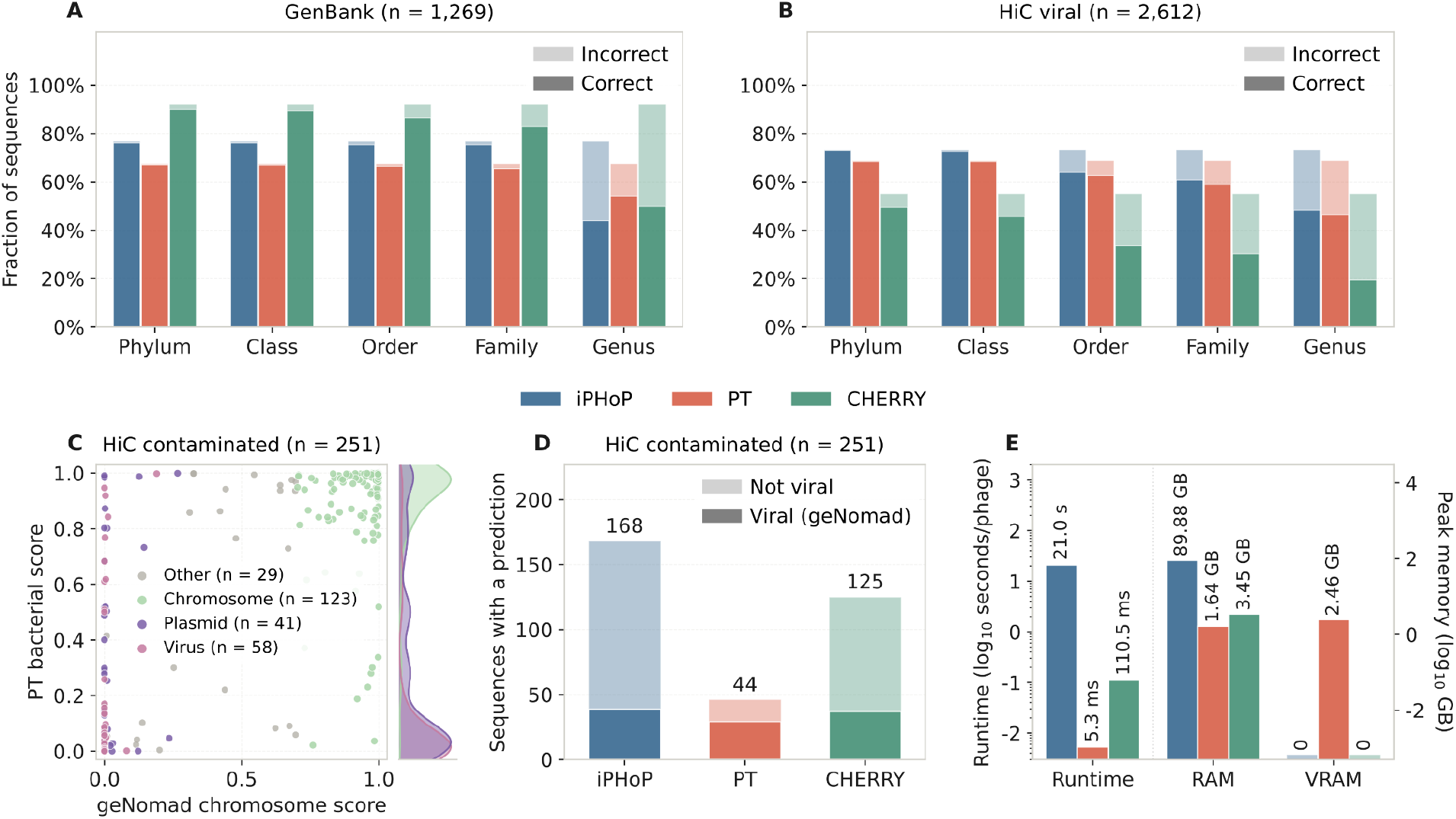
Comparison of PT with iPHoP and CHERRY on independent test sets. Host prediction accuracy across taxonomic ranks on **(A)** the GenBank dataset (n=1,269) and **(B)** the HiC viral dataset (n=2,612). **(C)** Comparison of PT’s bacterial fragment score with geNomad’s chromosome score on sequences in the HiC contaminated dataset. **(D)** Number of sequences that receive a confident host prediction in the HiC contaminated dataset. For PT, only predictions with a bacterial score below 0.5 were considered. **(E)** Average time per phage genome evaluated using all phage genomes from the GenBank and HiC viral datasets (n=3,881) on a system with an AMD EPYC 9124 CPU and a NVIDIA H100 GPU for iPHoP (20 threads), PT (batch size 16, 1 thread, 1 GPU), and CHERRY (20 threads). Runtime was measured using the $SECOND variable around each tool call. Peak memory was measured as the job cgroup v1 memory.max_usage_in_bytes recorded by Slurm, and peak GPU memory as the maximum of nvidia-smi memory.used sampled at 2 Hz.

We then turned to the 251 contigs of the HiC contaminated dataset. Classified by geNomad, most of these were not viral: 123 sequences (49%) received a chromosome score above 0.7 and a further 41 (16%) were called plasmid, against only 58 (23%) called viral. PT’s bacterial fragment score agreed closely with this assignment (Figure 4C). This was then reflected in the number of confident host predictions each tool produced (Figure 4D): iPHoP returned a prediction for 168 of the 251 sequences (67%) and CHERRY for 125 (50%), whereas PT returned one for 44 (18%). For these predictions, 28 of PT’s 44 corresponding sequences geNomad independently classified as viral (64%), compared with 39 of 168 for iPHoP (23%) and 36 of 125 for CHERRY (29%).

Finally, because PT requires only a single forward pass through a neural network architecture, it is much cheaper to run compared to the other tools if the underlying system has a GPU installed (Figure 4E). PT processed a phage genome in 5.3 ms on a single CPU core and a GPU, compared with 110.5 ms for CHERRY and 21.0 s for iPHoP on 20 CPU cores, roughly a 20-fold and 4,000-fold difference, respectively. The gap in memory usage was comparable: peak memory usage was 1.64 GB for PT and 3.45 GB for CHERRY, against 89.88 GB for iPHoP. PT additionally used 2.46 GB of GPU memory.

## Discussion

Host prediction remains an important but challenging step in viromics research. Here, we demonstrated that a neural network architecture trained on raw nucleotide sequences can predict phage hosts at genus-level resolution with accuracy comparable to tools that integrate diverse phage-host signals, including sequence alignment, protein-level homology, and curated reference databases. By processing DNA as trinucleotides across all six reading frames, the PhageTransformer model captures compositional biases and the coding structure of phage genomes without requiring gene calling or annotation as a preprocessing step.

Inspection of the model’s internal representations revealed that this performance is connected to biologically meaningful features. The learned codon embeddings recover amino acid identity, purine content, and physicochemical properties of the encoded residues, all without explicit supervision. Similar observations have been made for CaLM, a codon-level protein language model whose learned embeddings capture amino acid identity and biochemical properties from codon tokens, and which outperforms far larger amino acid-based models on tasks such as species recognition^53^. At the genome level, the cross-frame attention mechanism implicitly identified coding regions by assigning high importance to reading frames that carry genes even in genomes utilizing alternative genetic codes, paralleling the emergent learning of gene structure reported for the Evo genomic foundation models^23,24^. Together with the observed importance of tail proteins for PT’s predictions, these findings suggest that the model did not merely memorize sequence motifs, but learned generalizable representations of the relationship between nucleotide composition, gene content, and host taxonomy.

Our analysis revealed that phage genomes carry sufficient shared signals with their hosts for the model to confidently predict the correct bacterial genus even when applied to purely bacterial DNA. Some of these shared signals are well established: translational selection drives phage genes toward host-preferred codons, particularly those encoding structural proteins, which are required in high multiplicity during assembly and therefore have to be expressed efficiently^33^. Temperate phages show somewhat stronger codon adaptation than lytic phages, consistent with their prolonged co-replication with the host chromosome^34^. Still, lytic phages exhibit substantial codon adaptation to their hosts, indicating that the shared signal is not restricted to integrated prophages but reflects a general property of phage-host co-evolution. By incorporating a dedicated bacterial detection class during training, PT leveraged this shared signal to simultaneously predict hosts and flag non-viral sequences.

This dual functionality makes PT particularly suited for metagenomic applications, where assembled contigs often include bacterial contamination, misassembled chimeras, or proviral sequences embedded in host chromosomes. Rather than requiring a separate filtering step with tools such as geNomad or CheckV, PT can flag suspect contigs directly during host prediction. Combined with its speed, this enables analysis of large-scale metagenomic datasets where millions of contigs need to be screened. We anticipate that this combination of host prediction, contamination detection, and taxonomic classification of bacterial fragments will be valuable for studies investigating phage-host dynamics, viral ecology, and microbiome composition at scale.

PT achieves competitive performance for genus-level host prediction based solely on phage genome sequences, without requiring protein annotation, database searches, or host genome input. This independence from host sequence at inference time comes at a cost in flexibility. iPHoP and CHERRY allow users to supply additional host genomes and obtain predictions against them. PT instead has a fixed set of genera it can predict, and extending it to a new genus requires retraining, which is implemented as part of the tool. Since PT learns primarily from host-labelled phage sequences, adding a genus requires phage examples associated with it rather than the host genome alone. Future work will address this limitation with a host genome encoder against which PT can score its predictions, allowing generalization to bacterial genomes from unseen genera and host prediction beyond a fixed set of genera.

A second limitation concerns resolution. PT predicts hosts at the genus level, and we do not expect this architecture to extend straightforwardly to strain-level resolution, where phage-host specificity is governed by fine-grained molecular determinants such as receptor-binding protein-receptor compatibility and surface polysaccharide variation^5,54–56^. Recent work has demonstrated that strain-level prediction is feasible within individual host genera but requires genus-specific training data and mechanistic features that are not captured by genome-wide compositional signals alone^54–56^. Along similar lines, a recent review of AI-driven strain-level prediction approaches concludes that hybrid, modular systems combining broad-scope learned representations with genus-specific mechanistic priors offer the most credible path toward clinically useful resolution^57^. Future work could therefore combine a genus-level prediction model such as PT with an ensemble of smaller, genus-specific models that capture the fine-grained molecular determinants of infection.

Adversely, current understanding of phage-host interactions is still limited across a large number of poorly explored host genera: training data of any kind remains scarce outside a small set of model systems, and the genera for which fine-grained mechanistic determinants are known are largely the same ones for which cultured phage-host pairs already exist. Our training dataset substantially extends the number of host genera and the corresponding number of phage genomes over the training data used in other host prediction frameworks (Table 1). However, a large fraction of host genera is only represented by a handful of example genomes and we do not expect PT to generalize well in these, limiting its applicability in viromes derived from biomes other than human-associated ones. One challenge towards broadly applicable host prediction is therefore to scale up high-throughput methods for establishing phage-host links, and to make their output precise enough to train on.

A number of experimental approaches now generate phage-host associations without requiring either partner to be cultured, and each captures a different stage of the infection cycle. Viral tagging labels phage particles fluorescently and sorts host cells that have acquired a signal, linking phages to hosts at the point of attachment^58^. Adsorption sequencing (AdsorpSeq) instead separates phages bound to isolated cell envelopes by electrophoresis and sequences the bound fraction, additionally capturing phage receptor binding strength^59^. Proximity ligation methods such as Hi-C crosslink DNA molecules that are physically co-located, so that phage genomes replicating inside a cell become linked to the host chromosome^48,60^. Single-cell genomics or transcriptomics finally sequence individual sorted cells, recovering whatever viral sequence they contain^61^. All these methods contain inherent sources of noise that can limit how useful the resulting data can be for predictive machine learning.

Equally challenging, and as important as phage-host prediction itself, is developing a meaningful, independent benchmark to compare tool performance. Phage-host interaction data are scarce and of variable quality. Traditionally, experimental data are considered the most reliable, but this information is scattered in the literature, and many studies that performed large phage-host screens lack phage sequence data, especially older ones (for example, see references in Weitz et al.^8^). Many of the phage-host pairs that are present in public databases^35,38,40^ are already used to train the tools, so using them for benchmarking will lead to overlap between training and testing data. An additional reliable source is integrated prophages, which we used here, but identifying prophages is not trivial and could lead to overlap in sequence signal, as we also observed. We hope that the field will continue to search for innovative methods to measure phage-host interactions, especially those producing high-throughput data, which will form an important resource to benchmark the computational tools, and serve as training data for future predictive models.

## Data availability

Supplementary tables, model weights, the phage genome dataset and accession numbers of the host genome dataset were deposited at Zenodo in a combined archive (https://doi.org/10.5281/zenodo.22112416). This archive also contains the scripts used for analysis and visualization of the presented data. The trained PhageTransformer production model is available as a tool on github: https://github.com/MGXlab/phagetransformer

## Acknowledgements

We thank Yasas Wijesekara for valuable discussions on encoding sequence information with neural networks. This study was supported by the European Research Council (ERC) Consolidator grant 865694: DiversiPHI, and the Deutsche Forschungsgemeinschaft (DFG, German Research Foundation) under Germany’s Excellence Strategy – EXC 2051 – Project-ID 390713860, and the Alexander von Humboldt Foundation in the context of an Alexander von Humboldt-Professorship founded by the German Federal Ministry of Education and Research.

