## Supplementary Information for "PhageTransformer for scalable and accurate bacteriophage host predictions"

### PhageTransformer - scalable and accurate host assignments for bacteriophages

Malte Siemers[1], José L. López[1], Bas E. Dutilh[1, 2]
[1]Institute of Biodiversity, Ecology, and Evolution, Faculty of Biological Sciences, Cluster of Excellence Balance of the Microverse, Friedrich Schiller University Jena, Jena 07745, Germany
[2]Theoretical Biology and Bioinformatics, Science4Life, Utrecht University, Utrecht 3584 CH, The Netherlands

#### Supplementary Information

### Supplementary note 1

**Selection of architecture hyperparameters**

Architecture hyperparameters were selected by successively varying one quantity at a time and evaluating classification performance on the validation set, with training and inference cost treated as a secondary criterion. Several choices additionally follow from constraints imposed by six-frame tokenization and by the compression factor of the convolutional tower:

**Patch length and stride**

The patch length of 3,074 nt follows from a tokenization constraint. A patch of length $L$ tokenized in reading frame $f \in \{0, 1, 2\}$ yields $\lfloor (L - f)/3 \rfloor$ tokens, so the three frames of a given strand contain equal numbers of tokens only when $L \bmod 3 = 2$. At $L$=3,072, frame +1 would contain 1,024 tokens and frames +2 and +3 only 1,023, requiring padding of the shorter frames and introducing a systematic offset between frames at the patch boundary. Adding two nucleotides removes this asymmetry: at $L$=3,074 all six frames yield exactly 1,024 tokens, so the frames are aligned by construction and cross-frame attention operates on positions that correspond across frames without padding. The value 1,024 was chosen because it is divisible by the total compression factor of the convolutional tower (16, see below), so that the patch is compressed to a whole number of positions (64) with no remainder and no padding at the final convolutional layer. Patch lengths below 2,000 nt were tested and gave reduced classification performance, consistent with shorter patches truncating a larger fraction of genes and providing less context per patch. The stride of 2,400 nt produces an overlap of 674 nt between consecutive patches. This value was chosen with two considerations in mind. First, the receptive field of the convolutional tower spans 97 codon tokens, corresponding to 291 nt of sequence. An overlap of 674 nt exceeds twice the receptive field, ensuring that every position affected by a boundary in one patch is represented with full convolutional context in the neighbouring patch. Second, the overlap covers a substantial fraction of a typical bacterial gene, so that coding regions spanning a patch boundary remain largely intact in at least one patch.

**Codon embedding dimension**

The codon embedding dimension was increased from 4 to 32 to 128. Raising the dimension from 4 to 32 improved classification performance, whereas the increase from 32 to 128 did not. We nonetheless retained 128 dimensions based on the structure of the learned embeddings: principal component analysis of the 128-dimensional embedding matrix showed that PC30 still carried half the variance of PC1 (Figure S2), indicating that the learned codon representations occupy a substantially higher-dimensional subspace than 32 dimensions permits. We interpret the absence of a measurable performance gain as a limitation of the downstream capacity rather than evidence that the additional dimensions are unused, and kept the larger embedding dimension as the less constraining choice at negligible parameter cost.

##### Convolutional tower

The tower comprises four layers with kernel sizes 9, 9, 7, 7 and stride 2 throughout, giving 16-fold compression: the 1,024 tokens of each frame are reduced to 64 positions before the patch-level transformer. The compression factor was selected by comparing three configurations. 8-fold compression (128 positions entering the transformer) did not improve classification performance sufficiently to justify its higher memory footprint and slower training and inference, given the quadratic scaling of self-attention in sequence length. 32-fold compression (32 positions) reduced performance, indicating a loss of information at that resolution. Sixteen-fold compression was retained as a compromise between computational cost and accuracy. The channel dimension was increased from 300 to 600 in steps of 100. Performance improved up to 500 channels; the increase from 500 to 600 yielded only a marginal gain. We selected 600 and did not test larger values, as the returns were already diminishing relative to the associated increase in training and inference time.

##### Second convolutional branch

The second convolutional branch, which retains frame-resolved representations through an additional convolutional stage before merging, was introduced together with bacterial sequences for contamination detection. Its addition slightly improved the bacterial fragment detection rate. Increasing the number of layers (3) or filters (64) within this branch produced no further improvement, so the branch was kept at its minimal effective configuration.

##### Transformers, cross-frame attention, and pooling

Layer counts and model dimensions for both the patch-level and the contig-level transformer were selected by successive increase, and fixed at the point where further increases produced only minor gains in classification performance at a marked cost in training and inference time. Cross-frame attention and attention pooling both use a single learned query rather than multiple queries. A multi-query variant was tested and gave no improvement in classification performance while increasing training and inference time, so the single-query form was retained.

##### Loss function and output activation

Outputs are produced by independent sigmoid activations over the raw logits rather than a softmax, so that a contig may be assigned to several host genera simultaneously, to the bacterial fragment class concurrently with a genus, or to no class at all. We compared several loss functions compatible with multi-label prediction: binary cross-entropy, hill loss, and focal loss. Focal loss was selected because it increased recall for host genera with few training examples relative to the alternatives, consistent with its down-weighting of well-classified examples shifting gradient toward the rare and difficult classes that dominate the tail of the genus distribution in the course of the training.

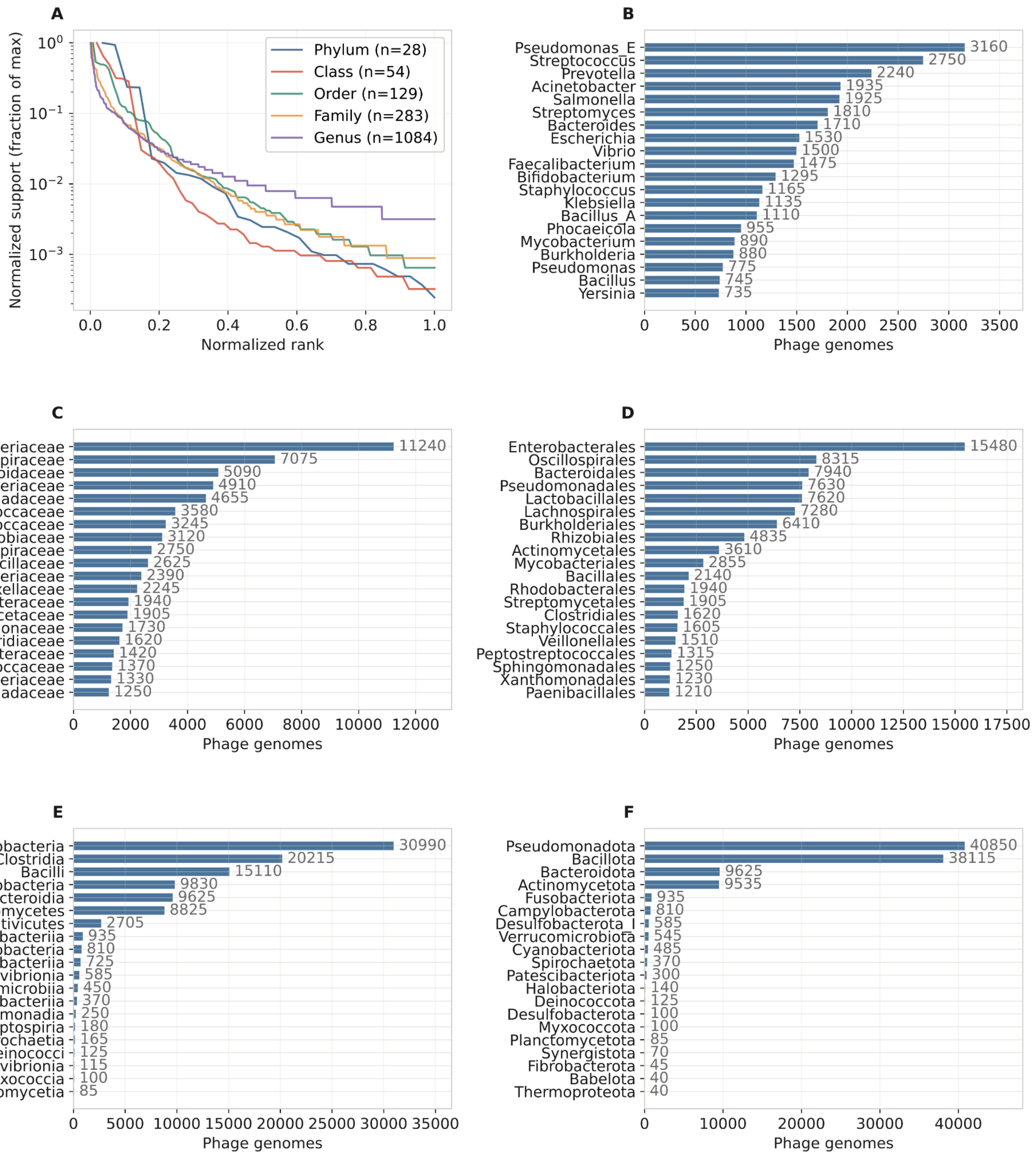


**Figure S1. Training dataset overview. (A)** Normalized distribution of the number of phage genomes across host taxonomic ranks. The dataset spans phage genomes with annotated bacterial hosts from 28 phyla, 54 classes, 129 orders, 283 families, and 1,084 genera. **(B-F)** The phages infect a variety of hosts. To show the skew in host taxa distribution, the bar plots show the 20 host taxa with the largest number of associated training viral genomes for taxonomic ranks genus **(B)**, family **(C)**, order **(D)**, class **(E)**, and phylum **(F)**.

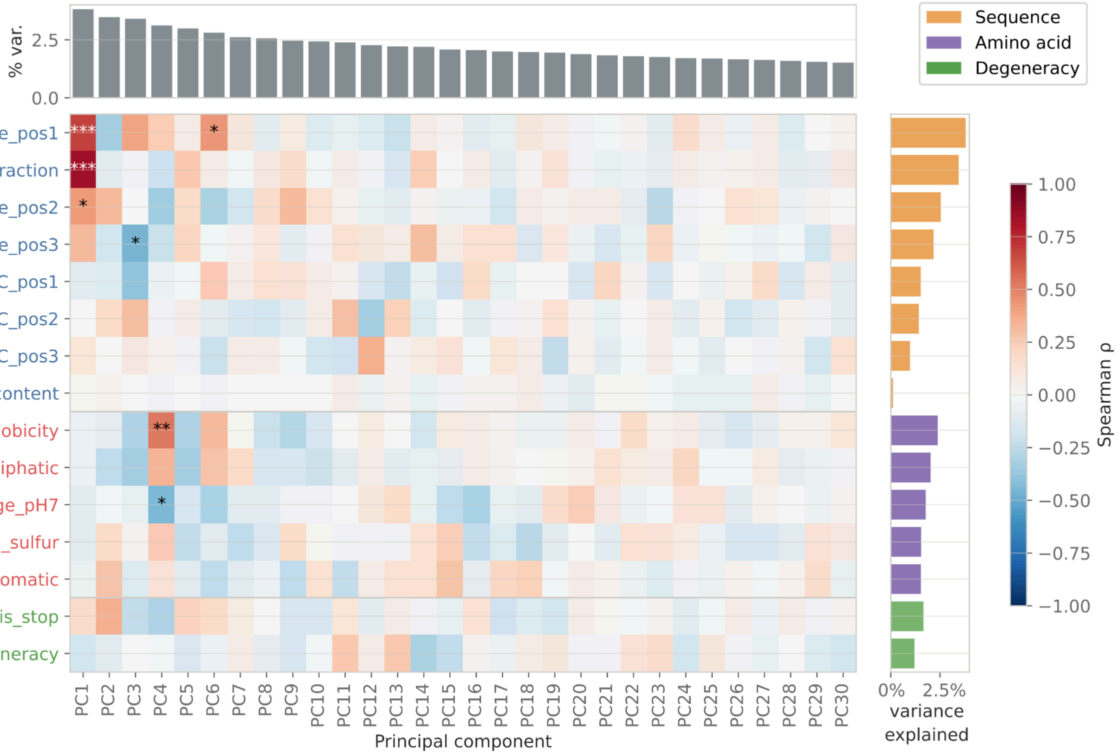


**Figure S2. Codon property–embedding correlation analysis.** Spearman rank correlations between 15 curated codon properties and the first 30 principal components of the learned codon embeddings. The top panel shows the percentage of variance explained by each PC. The right panel shows the total embedding variance explained by each property (variance-weighted $R^2$ summed across all PCs). Properties are color-coded by category (blue: sequence composition, red: amino acid property, green: degeneracy). Significance stars indicate BH-corrected q-values (* q<0.05, ** q<0.01, *** q<0.001).

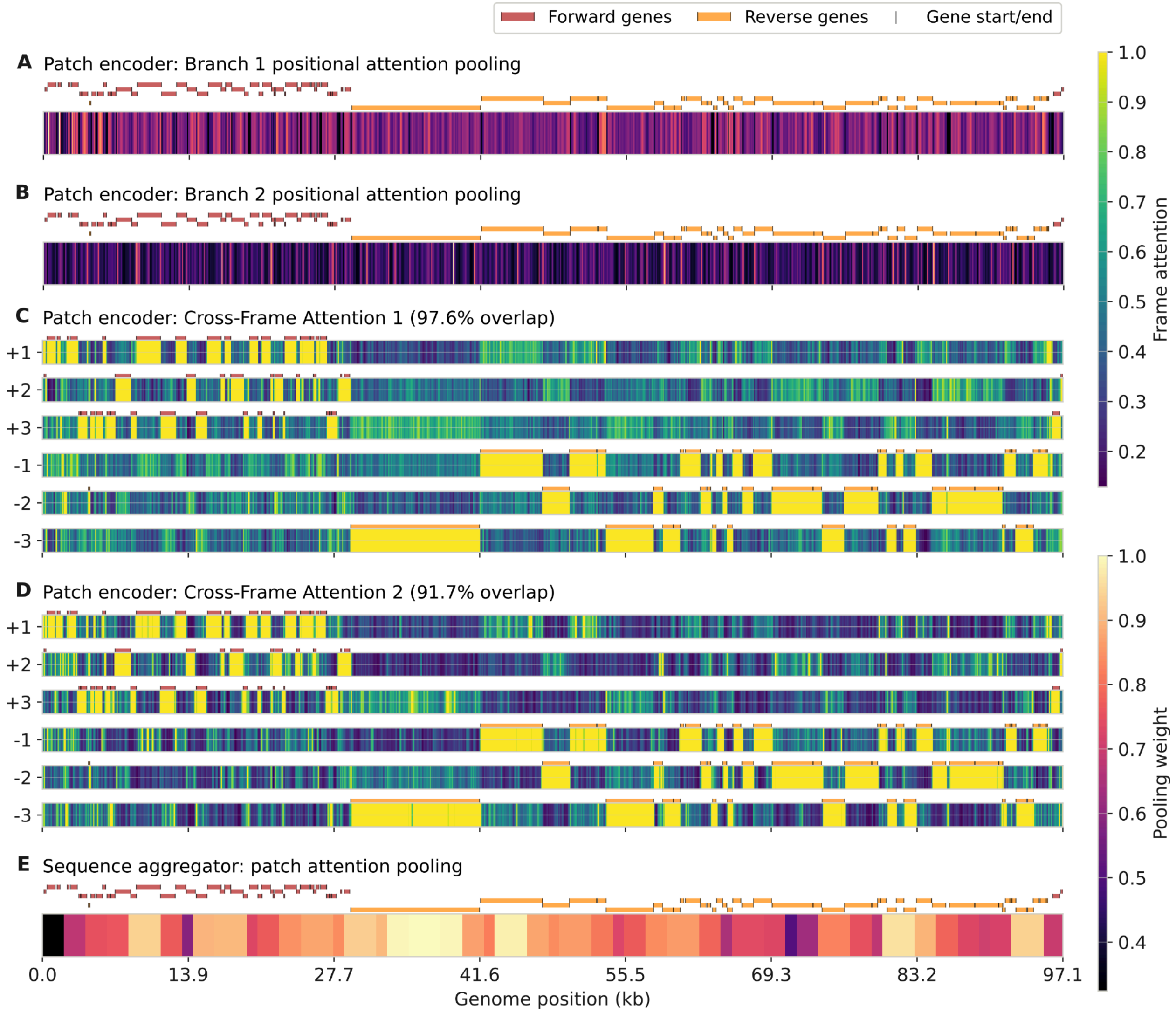


**Figure S3. Attention profiles across attention mechanisms along a single genome.** Positional attention pooling (AP in Figure 1) weights for **(A)** Branch 1 and **(B)** Branch 2 in the patch encoder, normalized attention weights for frame selection in the patch encoder for **(C)** Cross-Frame Attention 1, and **(D)** the Cross-Frame Attention 2 (see Figure 1), and **(E)** the patch attention weights in the sequence aggregator transformer along the genome of *Carjivirus communis* (BK010471.1, 97.1 kb). Gene locations, reading frames, and directions are indicated by colored strips on top of the attention heatmaps (red for forward, orange for reverse, black lines indicate gene boundaries). Higher attention values indicate a larger contribution to the final prediction.

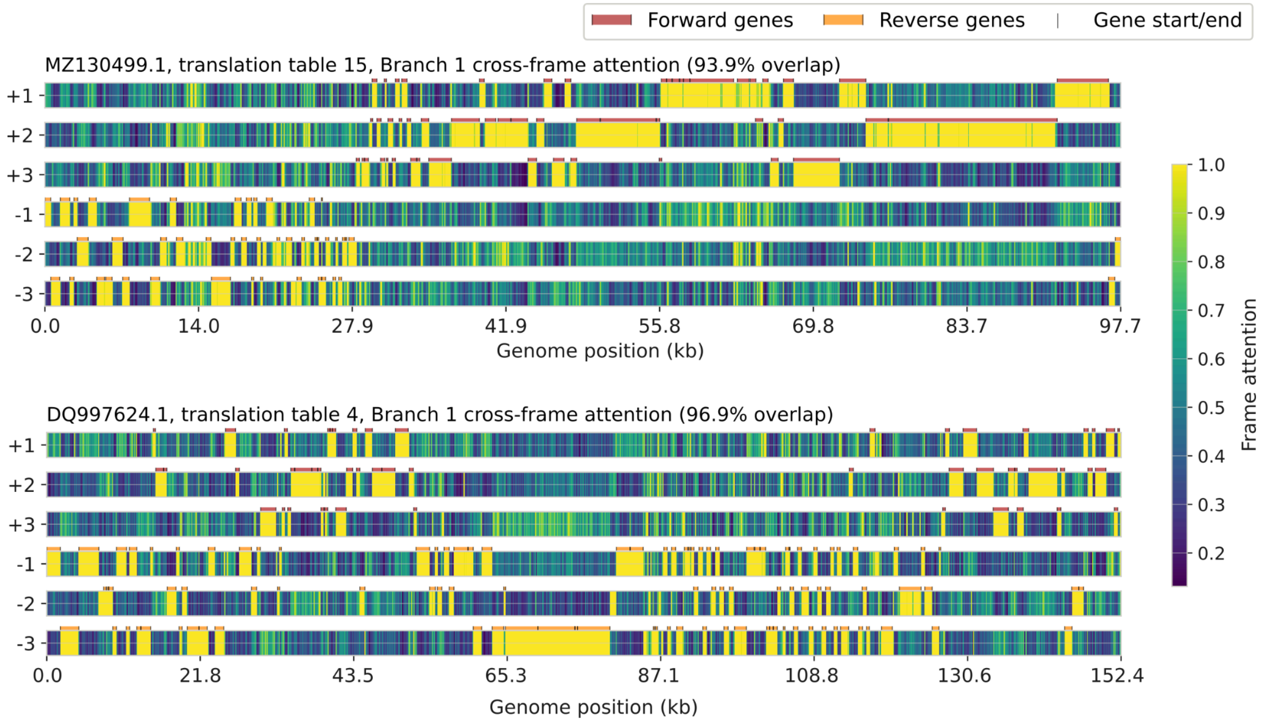


**Figure S4. Cross-frame attention weights for phages utilizing alternative genetic codes.** We extracted the attention weights of the patch encoder Cross-frame Attention 1 layer (see Figures 1 and S3 for more details) for two phage genomes and annotated CDS with pyrodigal-gv for *Cervivirus coli* with translation table 15 (MZ130499.1, top) and Thermus thermophilus phage YS40 with translation table 4 (DQ997624.1, bottom). Gene locations, reading frames, and directions are indicated by colored strips on top of the attention heatmaps (red for forward, orange for reverse, black lines indicate gene boundaries). Higher attention values indicate a larger contribution to the final prediction.

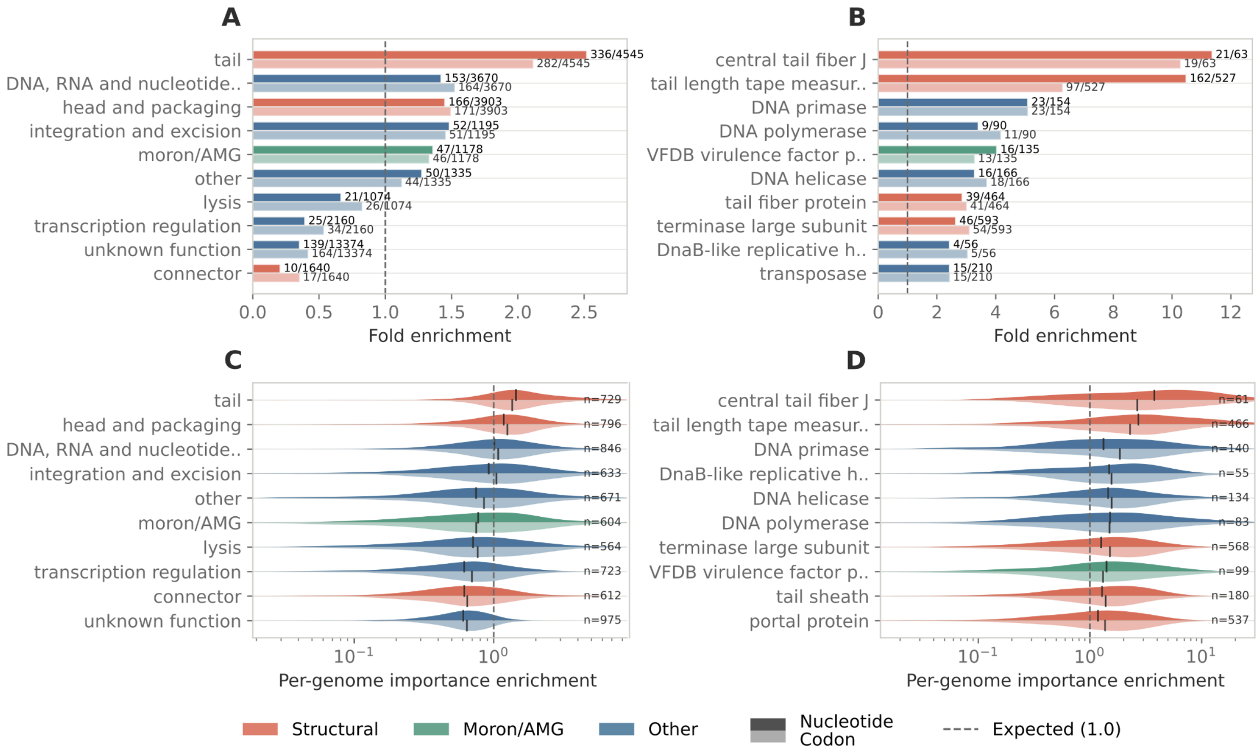


**Figure S5. Importance of PHROG categories and products towards host predictions.** We randomly sampled 1,000 phage genomes, 10 per host genus across the 100 genera with the most training samples, from the phage genome validation set. Pharokka (v1.9.1, DB version 1.8.0, CDS called with phanotate) and phold (v1.2.2) were used to annotate CDS with PHROG categories and products. In each genome, within the boundaries of each CDS, sequences were scrambled at the nucleotide level and at the codon level. PT was used to compute host scores for unmodified and per-CDS scrambled sequences. Importance per CDS was defined as the host score of the correct host genus for the unmodified version minus the score derived from the genome with the nucleotides or codons of the corresponding CDS scrambled. Fold-enrichment of an annotation was computed as the ratio between the fraction of genomes carrying this annotation with maximum importance and the fraction of CDS with that annotation across all genomes. Importance per annotation and genome was defined as the sum of the importance values for all CDS in one genome annotated with the respective category. Per-genome Importance enrichment of an annotation was computed as the ratio of importance of that annotation divided by the fraction of CDS carrying that annotation per genome. **(A)** Fold-enrichment of PHROG categories. **(B)** Top-10 fold-enriched PHROG products. **(C)** Distribution of per-genome importance enrichment for PHROG categories. **(D)** Distribution of per-genome importance enrichment for top-10 enriched PHROG products.

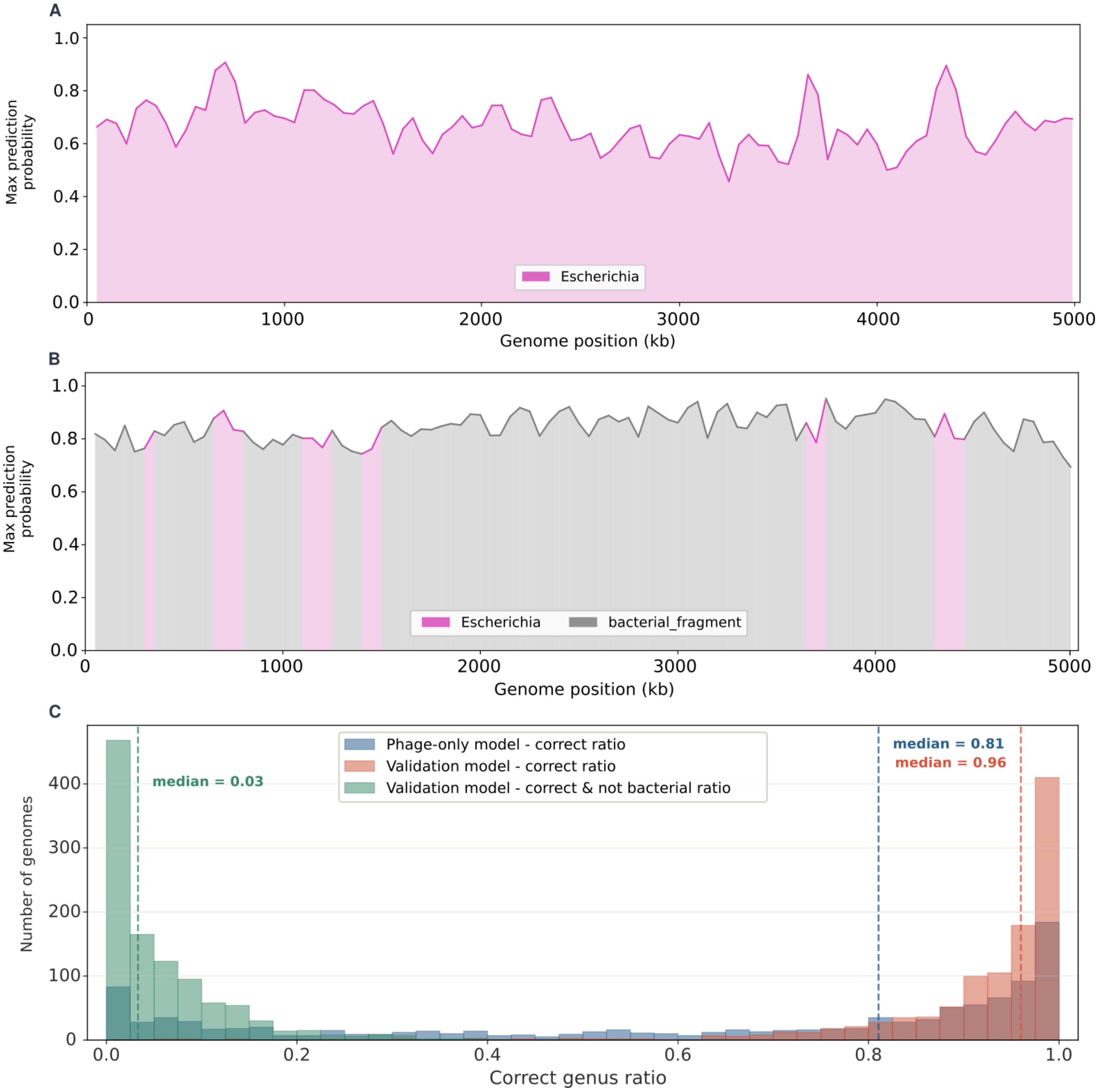


**Figure S6. Bacterial detection and taxonomic classification.** Comparison of classification scores along bacterial genomes between two model configurations, the phage-only model and the validation model (see Methods). Classification scores for the highest ranking host genus along an *E. coli* genome using **(A)** the phage-only model and **(B)** the validation model. **(C)** Distribution of correct classification ratios across 1,084 bacterial genomes randomly selected per host genus for both models. The host genus was predicted in 50,000 nucleotide long windows along the validation region in each genome and the ratio of correctly classified windows was recorded. Dashed lines indicate medians. For the phage-only model (blue, median 0.81), the highest-ranking genus was compared to the taxonomy of the bacterial genome. For the validation model, two distributions are reported: First, based on the highest-ranking genus prediction ignoring the bacterial score (red, median 0.96) and second, only matching windows with a bacterial score below 0.5 were considered correct (green, median 0.03).

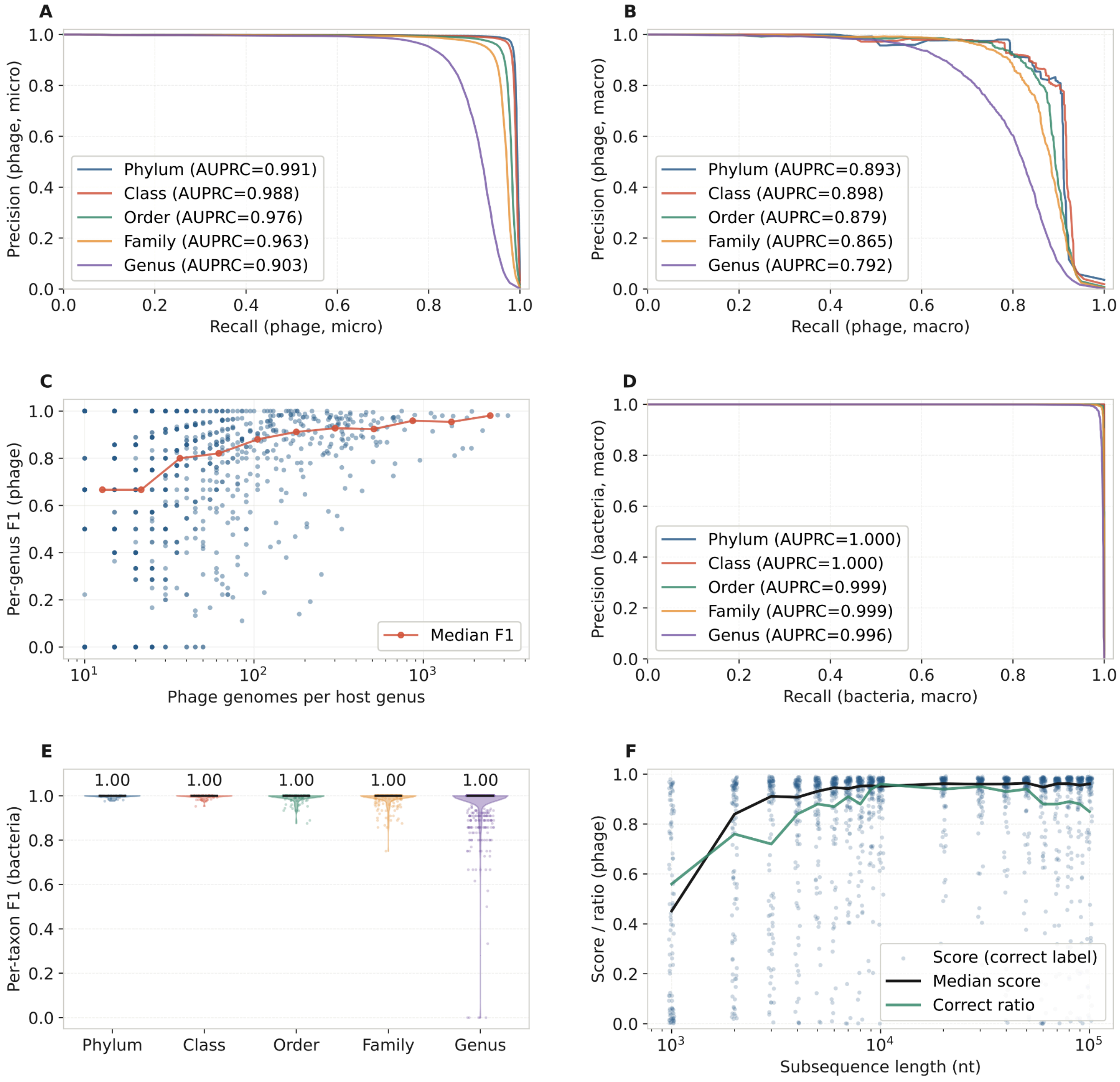


**Figure S7. Classification performance across taxonomic ranks.** Host-prediction metrics were computed on the held-out validation set of the phage genome dataset (20% of the full dataset, n=18,173 phage sequences) while taxonomic classification metrics were computed on sequences sampled from the validation regions in the 1,084 bacterial genomes selected for training the validation model. **(A)** micro-averaged precision-recall curves at each taxonomic rank **(B)** macro-averaged precision-recall curves at each taxonomic rank **(C)** Per-genus F1 score as a function of training support (number of phage genomes per host genus). Red line shows the binned median. **(D)** macro-averaged precision-recall curve for the taxonomic classification of bacterial sequences. **(E)** Violin plots of F1 scores per taxon at each taxonomic rank. Black lines and annotated numbers correspond to median F1 scores at each taxonomic rank. **(F)** Model confidence and correctness dependence on input sequence length (n=100 per reported length)

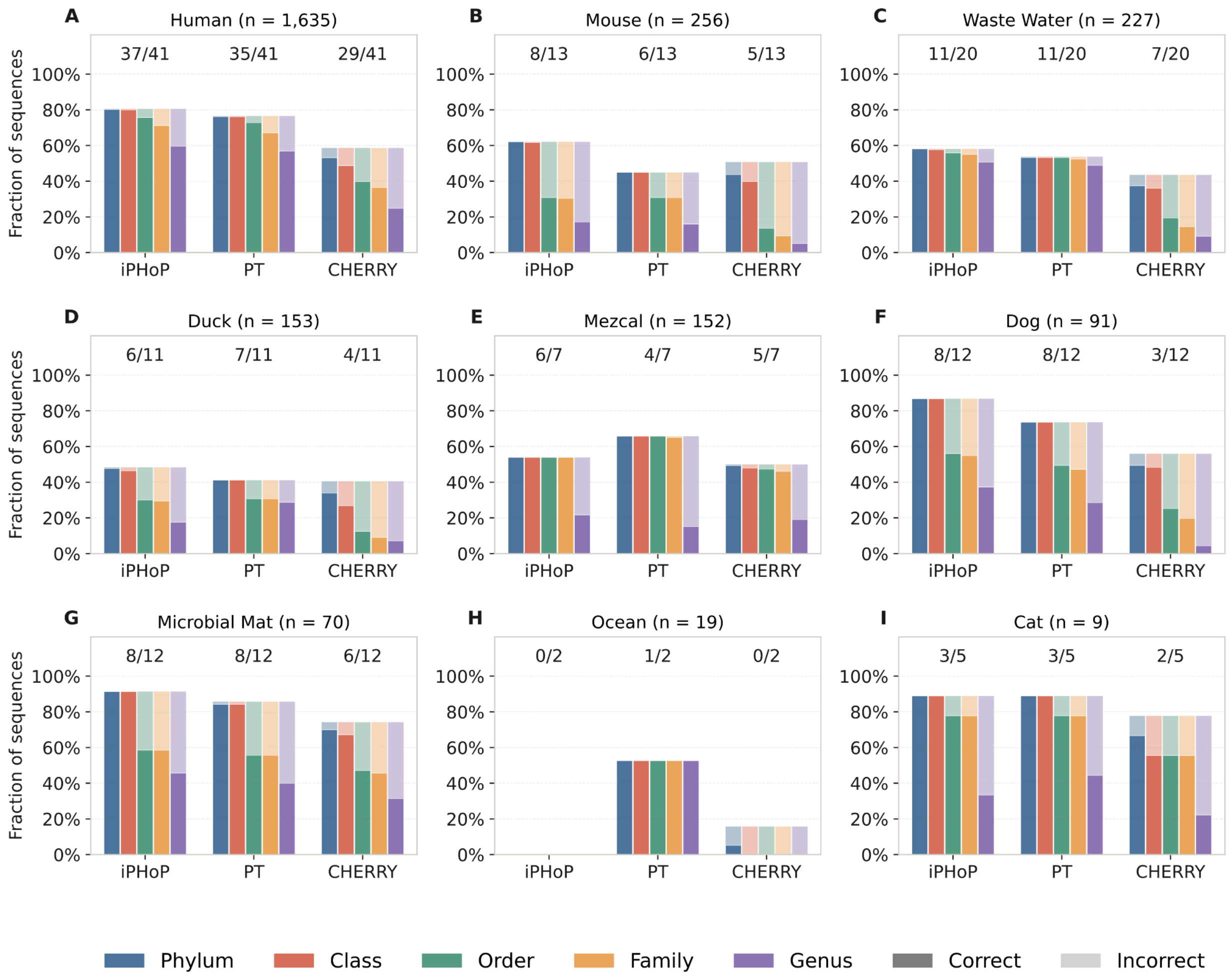


**Figure S8. Comparison of host prediction performance between PT, IPHoP and Cherry on the HIC viral dataset per biome.** Predictions of all three tools were recorded and together with the host genus labels translated to GTDB 226. For each tool at each taxonomic rank (phylum: blue, class: red, order: green, family: yellow, genus: purple), the fraction of sequences which received a prediction was split into correct (solid) and incorrect (transparent). In cases with multiple host labels or predictions per sequence, a sequence was considered correctly predicted if any of the predictions matched any of the labels. Prediction accuracy for sequences annotated with the biomes **(A)** human fecal, **(B)** mouse fecal, **(C)** waste water, **(D)** duck fecal, **(E)** mezcal fermentation, **(F)** dog fecal, **(G)** microbial mat, **(H)** ocean, and **(I)** cat fecal. Numbers on top of each group of bars indicate the number of host genera receiving at least one correct prediction out of all host genera present in the host labels.
